# Efforful verb recollection drives beta suppression in mesial frontal regions involved in action initiation

**DOI:** 10.1101/473355

**Authors:** Anna A. Pavlova, Anna V. Butorina, Anastasia Y. Nikolaeva, Andrey O. Prokofyev, Maxim A. Ulanov, Denis P. Bondarev, Tatiana A. Stroganova

## Abstract

Whether the motor cortex activation accompanying concrete verbs comprehension is necessary for verbs conceptual processing is still a hotly debated topic in the literature. Answering this question, we examined to what extent the more difficult access to verb semantics requires an additional engagement of cortical motor system in verb generation task. Using power suppression of MEG beta oscillations (15-30 Hz) as an index of sensorimotor activation, we presented to our participants the noun cues which either were strongly associated with a single verb and prompted the fast and effortless verb retrieval, or were weakly associated with multiple verbs and were more difficult to respond to. A whole-brain analysis of beta suppression revealed that the only cortical regions sensitive to the difficulty of semantic access were the higher order motor areas on the medial and lateral surfaces of the frontal lobe. This differential activation of cortical motor system accompanied effortful verb retrieval and preceded the preparation of vocal response for more than 500 milliseconds. Since the mid-frontal frontal brain areas are involved in maintaining abstract representations of actions during their initiating and planning, we argue that our finding supports the view that motor associations contribute to retrieval of verb semantics.

## Introduction

A major claim of the embodied approach to cognition is that semantic/conceptual processing critically depends on the same neural systems that are directly involved in perception or execution of the relevant experience (e.g. Binder & Desai, 2011; Kemmerer, 2015; Pulvermüller & Fadiga, 2010). This hypothesis has inspired a multitude of neuroimaging studies examining the relationship between the cortical motor system and the conceptual processing of action-related language.

The fMRI research have demonstrated that processing of action verbs and sentences, indeed, engages the motor-related portion of frontal lobe, albeit with a great variability among the reported localizations of the language-induced effects. An initial claim for a somatotopically organized pattern of the action verb-induced activation in the left primary motor cortex (Hauk, Johnsrude, & Pulvermüller, 2004) was not replicated consistently (Tomasino, Werner, Weiss, & Fink, 2007; Willems, Toni, Hagoort, & Casasanto, 2010; Zhang, Sun, & Wang, 2018); and other works found the action language-related response in the left premotor cortex, including its ventral portion incorporated into Broca's complex (Aziz-Zadeh & Damasio, 2008; Tettamanti et al., 2005). Another fMRI study showed that the same left ventral premotor regions were activated by passive listening to meaningless monosyllables and attributed this activation to subvocal auditory-to-articulatory mapping (Wilson, Saygin, Sereno, & Iacoboni, 2004). Supporting this idea, the left ventral premotor region was shown to be equally activated by action verbs, non-action nouns and pseudowords (de Zubicaray, Postle, McMahon, Meredith, & Ashton, 2010). To further complicate the picture, Postle with colleagues (Postle, McMahon, Ashton, Meredith, & de Zubicaray, 2008) reported that the only cortical region, which was sensitive to action verbs compared to non-action lexical stimuli was located within the medial wall of the frontal cortex at the medial supplementary and pre-supplementary areas (SMA and pre-SMA respectively) implicated in action initiation (Nachev, Kennard, & Husain, 2008).

Despite the existing controversy, most of the fMRI studies on language-motor coupling agrees on the recruitment of higher-order areas of motor cortex by action language processing (Jirak, Menz, Buccino, Borghi, & Binkofski, 2010). Nonetheless, as the opponents of the embodied approach point out, the fMRI data alone are not sufficient to determine the role and even the timing of this motor system involvement (Caramazza, Anzellotti, Strnad, & Lingnau, 2014; Chatterjee, 2010; Mahon & Caramazza, 2008). The observed motor activation may not be necessary for language comprehension, but serves to enrich semantic processing after a word meaning has been successfully accessed.

The transcranial magnetic stimulation (TMS) studies, which are theoretically capable to establish a causal role of motor system in action language, have also provided inconsistent results. For instance, Pulvermüller, Hauk, Nikulin and Ilmoniemi (2005) observed that lexical decision on leg-related verbs was faster after a single-pulse TMS applied to the leg-site of left primary motor cortex than after both sham and arm-site stimulation. However, Tomasino, Fink, Sparing, Dafotakis and Weiss, (2008) reported that TMS stimulation of the primary motor hand area increased the speed of decision on hand-related verbs only if a task required imagining the associated action. This suggests that effector-specific regions of primary motor cortex rather participate in post-retrieval imagery than in semantic processing of action-related language per se. Notably, all the available single-pulse TMS studies applied TMS over primary motor cortex, which contrasted with the fMRI results implicated rather the higher-order motor circuitry in action-language processing.

Taken together, the available evidence does not provide the coherent picture on the role of the motor system in action-language semantics. Time-resolved neural evidence from magnetoencephalography (MEG) – a methodology combining high temporal and spatial resolution – would be beneficial for delineating the role of lower- and higher-order motor cortex in action language. Here, we aimed to use MEG for examining a prediction derived from the embodied cognition hypothesis. If the motor system involvement contributes to verbs’ semantic retrieval, then (i) motor activation should well precede the response required by semantic task, and (ii) the activation strength should be modulated by the difficulty of semantic retrieval.

To manipulate the semantic load we used a verb generation (VG) task (Thompson-Schill, D’Esposito, Aguirre, & Farah, 1997) which is a semantic association task where participants overtly produce related verbs in response to visually presented noun cues. The difficulty of semantic retrieval varies with the number of verbs associated with a particular noun cue: the nouns are either strongly associated (SA) with a single verb and, thus, are easy to respond to, or weakly associated (WA) with many appropriate verbs and require additional effort to find a related verb in memory (see for a discussion Martin & Cheng, 2006; Snyder & Munakata, 2008).

To measure the strength and timing of motor system activation we focused on the neural oscillations in the beta frequency band (15-30 Hz). Beta oscillations are generated in cortico-basal ganglia-thalamo-cortical loops and are inherently tied to the motor functioning (Engel & Fries, 2010; Gilbertson et al., 2005). They are prominent during postural maintenance and are suppressed (or desynchronized) during preparation, initiation and execution of new motor sequences (Leocani, Toro, Manganotti, Zhuang, & Hallett, 1997; Pfurtscheller, Graimann, Huggins, Levine, & Schuh, 2003; Zhang, Chen, Bressler, & Ding, 2008). Although beta suppression, or beta event-related desynchronization (ERD), is highly characteristic for activation of motor system, it can be also be observed in other cortical regions during the tasks requiring the integration of complex, information-rich neural representations with motor output (e.g. Leventhal et al., 2012). Thus, MEG beta suppression appears to be a well-justified measure to estimate the language–motor interplay in verb generation task.

The previous MEG studies of verb generation task examined the beta suppression mainly in the context of the expressive speech lateralization (Findlay et al., 2012; Fisher et al., 2008; Pang, Wang, Malone, Kadis, & Donner, 2011; Youssofzadeh, Williamson, & Kadis, 2017). The effects of semantic retrieval demands on the observed beta suppression, as well as its cortical topography and timing fell outside their scope.

To evaluate the role of motor circuits in verbs’ semantic processing, we investigated the temporal dynamics of sensorimotor beta oscillations accompanying the period of search for target verb in semantic memory in VG task. Considering that the verb search is a protracted process with uncertain and variable onset but which uniformly terminates with response production, we defined the search period in relation to response onset in contrast to the previous VG studies, which analyzed the stimulus-related neural activity. Since the involvement of lower- and higher-order motor areas in verb processing remains to be elucidated, we employed whole-brain analysis with a rigorous statistics to determine the cortical regions in which the beta ERD is sensitive to semantic load. On the assumption of the essential role of motor circuitry in verbs conceptual representation, we expected that effortful retrieval of the required verb would be accompanied by greater beta suppression in motor area/areas of frontal lobe.

## Method

### Participants

Thirty-five volunteers (age range 20–48 years, mean age 26 years, 16 females) took part in this study. The participants were native Russian speakers, right-handed, with normal or corrected-to-normal vision. All the subjects had either complete or incomplete higher education and reported no neurological diseases or dyslexia. One participant was subsequently excluded from the analysis due to extremely low number of correct responses and another one due to MEG acquisition error. The final sample comprised 33 subjects. A written informed consent was obtained from all subjects. The study was conducted according to the principles of the Declaration of Helsinki and was approved by the Ethics Committee of the Moscow State University of Psychology and Education.

### Materials

The stimuli list comprised 65 Russian nouns with a single strong verb associate and 65 nouns that were weakly associated with many verbs. The nouns were selected in an independent norming study (for details see Butorina et al., 2017). If the majority of the norming sample (from 58% to 90%) responded with the same verb to a presented noun, the noun was included into the Strong Association (SA) category (e.g., ‘‘solovey—poyet/nightingale—sings’’). If less than 23% of the participants agreed on the same response, the noun was assigned to the Weak Association (WA) category (e.g., ‘‘bumaga— goryt, mnetsya, rvetsya/paper—burns, becomes crumpled, is torn’’). The mean word length, word frequency and number of lexical associates were matched between the SA and WA categories (Table 1). Word frequency was taken from Lyashevskaya and Sharov (2009) frequency dictionary; a number of lexical associates were taken from Russian Associative Thesaurus (Karaulov et al., 2002).

**Table 1.** Means (and standard deviations) for psycholinguistic parameters of the nouns having a single strong verb associate (SA) and those having multiple weakly associated verbs (WA).

|  | SA | WA |
| --- | --- | --- |
| Length in letters | 5.6 ($\pm 1.6$) | 5.7 ($\pm 1.45$) |
| Word form frequency (occurrence per million) | 50.6 ($\pm 87.3$) | 49.1 ($\pm 65.8$) |
| Number of lexical associates* | 72.9 ($\pm 52.6$) | 85.6 ($\pm 59$) |

\* irrespective of associates' grammatical class

### Design

The noun cues were presented in a white font on a black background on a screen placed at 1.5 m in front of the participant. The size of the stimuli did not exceed 5^°^ of visual angle. The participants were required to overtly produce a verb associated with a presented noun by answering a question: “What this noun does?”. The subjects performed the task continuously within 16 blocks, each contained eight stimuli of the same category in the random order. The noun cues appeared on the screen for 3500 ms and was preceded by a white fixation cross presented for 300–500 ms (Figure 1A). Blocks contained SA and WA nouns were alternated with 16 sec interval. The experiment was implemented in the Presentation software (Neurobehavioral Systems, Inc., Albany, CA, USA).

**Figure 1.**
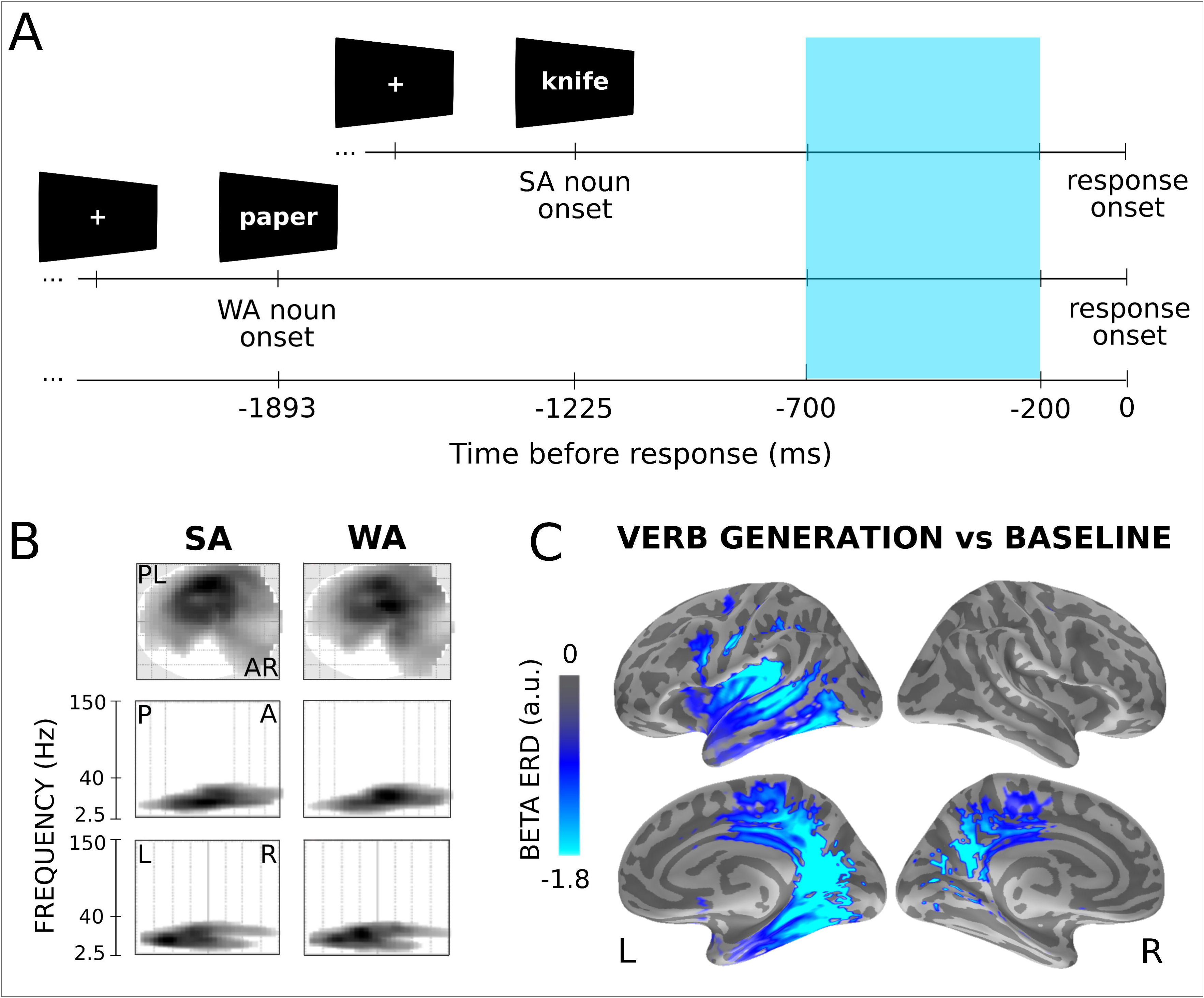
(A) Verb generation task design. The subjects were required to name a verb to the noun cue, which had either a single strong verb associate (SA) or many weak ones (WA). Time scale is aligned relative to the vocal response onset. Position of SA and WA nouns onsets is defined by mean RT for the respective trials. The blue rectangle represents the time window of interest used in the subsequent analysis. (B) SPM space-frequency statistical maps show scalp topography and frequency range of MEG power suppression (p < 0.05, FWE-corrected). A – anterior, P – posterior, L – left and R – right parts of the sensor array. (C) Reconstructed cortical sources of beta suppression (15-30 Hz) for SA and WA trials pooled together. Colorbar represents the strength of beta suppression in arbitrary units. The colored areas indicate the cortical regions with significant differences from baseline (p<0.05, Bonferroni-corr.).

### Behavioral responses

In course of verb generation session, participants’ vocal responses were tape recorded and off-line checked for response errors. The trials with no or semantically unrelated responses, grammatical mistakes, incomprehensible verbalization, imprecise vocalization onsets, pre-stimulus intervals overlapped with the vocal response to the previous stimulus were excluded from the subsequent analysis.

The onsets of participants' overt response were registered by the three-axis accelerometer placed on the participant’s throat (ADXL330 iMEMS Accelerometer, Analog Devices, Norwood, MA, USA). Speech onsets were marked using an automated algorithm (Zakharova et al., 2012) that detected increases in the accelerometer signal (z axis) above baseline by three standard deviations within a 3500 ms post-stimulus time window, and then visually inspected for false positives. The resulting reaction times and error rates were subjected to the repeated-measures analysis of variance (rmANOVA).

### MEG Data Acquisition

MEG data were acquired inside a magnetically shielded room (AK3b, Vacuumschmelze GmbH, Hanau, Germany), using a dc-SQUID Neuromag Vector View system (Elekta-Neuromag, Helsinki, Finland) with 204 planar gradiometers and 102 magnetometers. Data were sampled at 1000 Hz and filtered with a band-passed 0.03–333 Hz filter. The participants’ head shapes were measured by a 3Space Isotrack II System (Fastrak Polhemus, Colchester, VA, USA) by digitizing three anatomical landmark points (nasion, left and right preauricular points) and additional randomly distributed points on the scalp.

While recording, the position and orientation of the head were monitored by four Head Position Indicator coils. The electrooculogram was registered with two pairs of electrodes located above and below the left eye and at the outer canthi of both eyes for recording of vertical and horizontal eye movements respectively.

After MEG data acquisition, 28 participants underwent MRI scanning with a 1.5 T Philips Intera system to obtain a head geometry for later reconstruction of the cortical surface.

### MEG Pre-Processing

The raw data were subjected to the temporal signal space separation (tSSS) method (Taulu, Simola, & Kajola, 2005), embedded in MaxFilter program (Elekta Neuromag software), aimed to suppress magnetic interference coming from sources distant to the sensor array. Biological artifacts (cardiac fields, eye movements, myogenic activity), were corrected using the SSP algorithm implemented in Brainstorm software (Tadel, Baillet, Mosher, Pantazis, & Leahy, 2011). To compensate for the within-block head-movement (as measured by Head Position Indicator coils) a movement compensation procedure was applied. For sensor-space analysis, the data were converted to standard head position (x = 0 mm; y = 0 mm; z = 45 mm) across all blocks.

Data were divided into epochs of 1500 ms, comprising 500 ms before the stimulus presentation and 1000 ms before the vocal response onset. Epochs were rejected if the peak-to-peak value over the epoch exceeds 3*10^-10 T/m (gradiometers) and 12*10^-10 T/m (magnetometers) channels. Average number of verb generation trials finally taken for the analysis was 63+/−2 in SA condition and 53+/−5 in WA.

### MEG Data Analysis

Sensor-space data were analyzed using Matlab toolboxes SPM12 (Statistical Parametric Mapping software, Wellcome Trust Centre for Neuroimaging, London). For time-frequency analysis, we used a multitaper power spectrum estimation (Thomson, 2000), implemented in SPM12. We estimated the spectra in overlapping windows of 400 ms (shifted by 50 ms) with the frequency resolution set to 2.5 Hz for the frequency range 1-25 Hz, to 0.1*frequency for 25–50 Hz, and then to a constant 5 Hz resolution. These settings resulted in a single taper being used for 2.5–30 Hz, two tapers for 32.5–42.5 Hz, and three tapers for 45 Hz and above. The epoched time–frequency images were averaged over trials using a robust averaging procedure (Holland & Welsch, 1977). To normalize power changes across different frequency bands, the averaged power was log transformed and baseline corrected using a period of 300 to 0 ms before noun cue onset as the baseline (LogR option in SPM). Planar channels were then combined by adding time-frequency data for pairs of channels corresponding to orthogonal sensors at the same location.

We chose the period from −700 to −200 ms before the vocal response as time window of interest to study the response-related suppression. Considering a difference in the response times between SA and WA conditions (1.22 and 1.89 sec respectively), the chosen time window is well suited for between-condition comparison of verb retrieval period. Its lower boundary allows to omit an early time interval devoted to noun cue processing (≈500-600 ms after cue onset) even for SA condition, while the upper boundary excludes articulation preparatory period within approximately 200 ms before a vocal response onset (Bouchard, Mesgarani, Johnson, & Chang, 2013; Brumberg et al., 2016).

To define the frequency boundaries of beta suppression, we performed the statistical analysis of the response-related spectra in each experimental condition (SA and WA) to detect significant differences from the baseline. To this end, we averaged the time-frequency data over the period from −700 to −200 ms before the vocal response.

The 3D files of space (32*32 pixels) and frequency (60) dimensions were converted to Neuroimaging Informatics Technology Initiative (NIfTI) format and smoothed using a Gaussian smoothing kernel with Full Width Half Maximum of 8 mm * 8 mm * 3 Hz to ensure that the images conform to the assumptions of Random Field Theory (Kilner & Friston, 2010). Then, the smoothed images were subjected to a single-sample t-test with the family-wise error (FWE) correction for multiple comparisons at the cluster-level threshold of p < 0.05.

The adjacent frequency bins which showed significant spectral power suppression under both experimental conditions were collapsed into a single band of interest, for which power time-courses were computed for both experimental conditions. To ensure that the power suppression within the band of interest occurred throughout the entire verb retrieval period and was not restricted to its later part which could overlap with motor preparation period, we subjected the scalp * time images under each condition to a single-sample t-test (FWE-corr. at the cluster-level threshold of p < 0.05).

The final step was to contrast the power suppression within the band of interest under SA and WA conditions against each other in space and time dimensions. The scalp * time images were subjected to paired t-test and the resulting spatial-temporal clusters of significant differences were FWE-corrected for multiple comparisons. At the sensors with maximal between-condition differences we reconstructed power time-courses of the frequency band of interest for both experimental conditions. The amplitudes of maximal power changes against baseline were subjected to Wilcoxon Signed Rank test (with false discovery rate (FDR) correction for multiple comparisons).

### MEG source localization

Individual structural MRIs were used to construct a single-layer boundary-element models of cortical gray matter with a watershed segmentation algorithm (FreeSurfer 4.3 software; Martinos Center for Biomedical Imaging). Induced power in the source space was computed using dynamic statistical parametric mapping (dSPM) localization method (Dale et al., 2000) and Morlet wavelets with 2 cycles per wavelet, implemented in MNE Python open-source software (http://www.nmr.mgh.harvard.edu/martinos). This method provided time-courses of the induced power changes within the selected frequency band at each cortical location for each participant in each experimental condition. The individual data were then morphed into a common cortical model to allow a group comparison.

In order to examine which cortical regions displayed the verb retrieval-related suppression in the selected frequency band the data from two experimental conditions were collapsed, averaged within −700 - −200 pre-response time window and, using paired t-test, contrasted with the pre-stimulus baseline. The results of the statistical test were adjusted by using Bonferroni's correction for multiple comparisons.

To localize the between condition effects, the spatiotemporal data from SA and WA conditions were contrasted against each other. We restricted our statistical analysis to those time windows that demonstrated significant SA-WA differences at the sensor level. Providing that the sensor-space results had been corrected for multiple comparisons, in the source space the statistical threshold was set to uncorrected p < 0.05 in combination with a cluster extent of 10 adjacent voxels.

## Results

### Behavioral results

Consistent with previous VG studies (e.g. Snyder, Banich, & Munakata, 2011; Thompson-Schill et al., 1997), responses were faster and more accurate when the noun cues had a single dominant verb association (M = 1.22 +/− 0.17 s; 1.76 +/− 2.28% errors) compared to when the cues were weakly associated with multiple verbs (M = 1.89 +/− 0.23 s, 11.76 +/− 5.51% errors). rmANOVA revealed a highly significant effect of association strength on reaction times (F(1,32) = 325; p < 0.0001; η^2^ = 0.91) and error rate (F(1,32) = 167.6, p < 0.0001; η^2^ = 0.84).

## MEG results

### Frequency boundaries and spatial-temporal features of response-related beta suppression

#### Sensor-level analysis

Since beta frequency range varies between different VG studies (Findlay et al., 2012; Fisher et al., 2008; Youssofzadeh et al., 2017), as the first step we explored the frequency boundaries for VG task-induced beta suppression under SA and WA conditions. To this end, the data in each experimental condition were averaged across the −700 - −200 ms time window preceding the onset of vocal response and compared to the respective baselines using scalp-frequency SPM analysis. This time interval was characterized by highly significant oscillatory power suppression, which frequency boundaries largely overlapped between SA and WA conditions: 15-25 Hz for SA and 18-30 Hz for WA (both p’s < 0.0001, FWE-corr.; see Figure 1B). The above frequency ranges generally comply with the definition of beta oscillations (Engel & Fries, 2010). The beta ERD was characterized by a widespread, predominantly left-lateralized scalp topography, which was also similar for SA and WA (see Figure 1B). For the subsequent analysis we chose beta frequency band of 15 - 30 Hz, which covered all frequencies that underwent significant power attenuation under either task condition.

Next, we examined the temporal dynamics of the beta ERD using space-time SPM analysis. The analysis revealed that beta oscillations were reliably suppressed at each consecutive 50 ms interval throughout the entire −700 - −200 time window preceding vocal response (p < 0.0001, FWE-corr.). The comparison of response-related and stimulus-related beta ERD time courses (Figure 2B) revealed that the selected pre-response period captured the maximum of gradually developing beta suppression, which did not attenuate until the vocal response onset.

**Figure 2.**
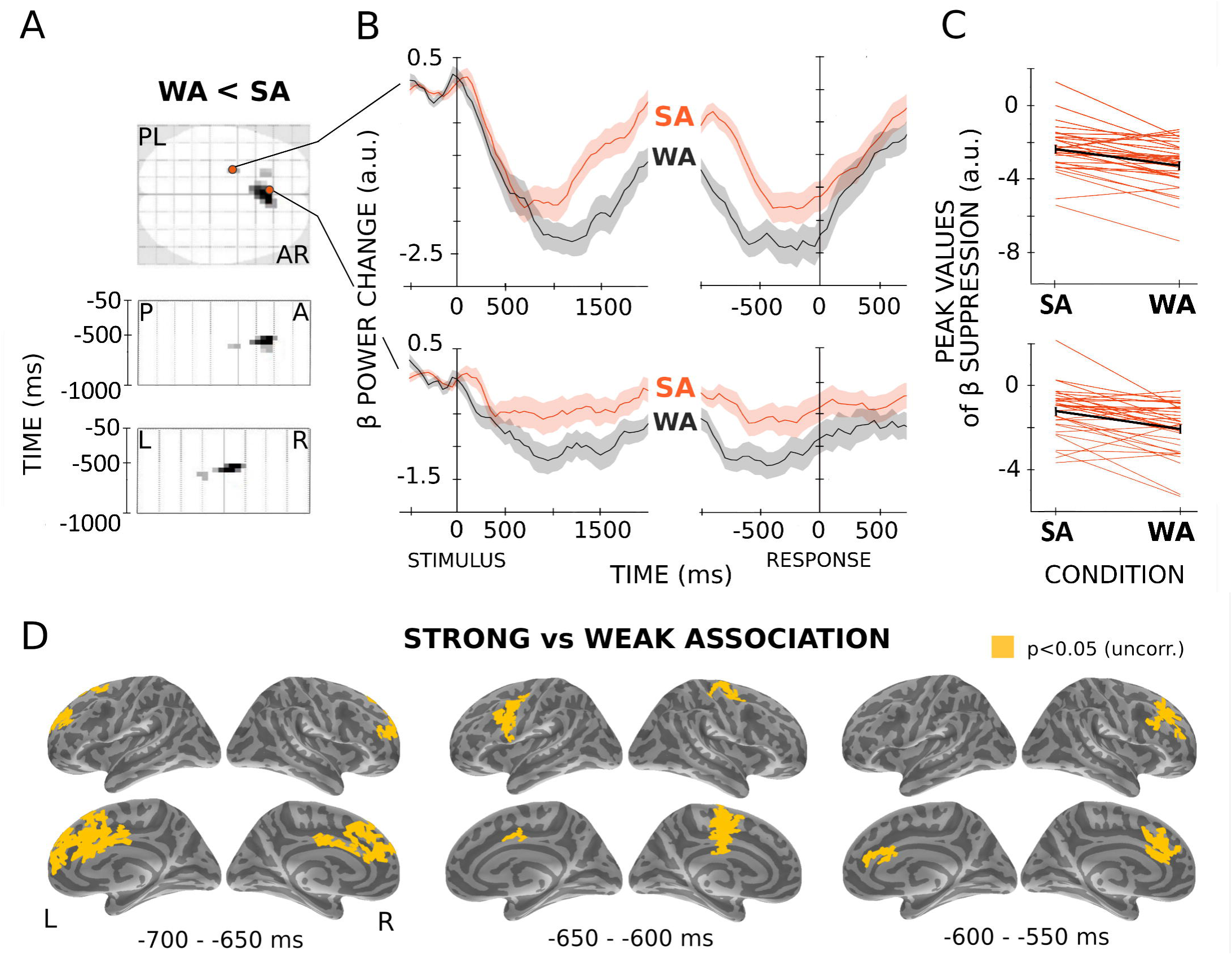
Differences in beta suppression between Strong (SA) and Weak Association (WA) trials. (A) Sensor-based SPM space-time images show three statistical clusters demonstrating significantly stronger response-related beta suppression in WA compared to SA trials (p < 0.05, FWE-corr.). A – anterior, P – posterior, L – left and R – right parts of the sensor array. (B) Time-courses of beta (β) power changes under SA and WA conditions calculated from the sensors nearest to maxima of two most significant clusters. The grand average time-courses are aligned to the onset of the noun cue (left panel) and to the onset of the vocal response (right panel). The shaded areas denote the standard error of the mean. Note, the stronger beta suppression corresponds to lower (more negative) values of beta power change. (C) The individual peak values of beta suppression at the same sensors under SA and WA conditions. The thick black line represents the group medians (Wilcoxon matched pairs test, p < 0.05). (D) The reconstruction of cortical sources corresponding to the three sensor-space temporospatial clusters. The images were thresholded at p<0.05 (uncorr.).

#### Source-level analysis

Figure 1C represented the cortical regions exhibiting the robust beta ERD under the VG task (p < 0.05, Bonferoni-corr.), the SA and WA conditions were pooled together for this analysis. In accordance with the previous findings (Findlay et al., 2012; Fisher et al., 2008; Pang et al., 2011; Youssofzadeh et al., 2017), the beta-band suppression was strongly lateralized to the left hemisphere, and comprised the widely distributed cortical network including the areas of temporal lobes involved in language processing as well as the regions on the medial and lateral surface of the frontal lobe subserving initiation, planning and execution of motor programs (see Table 2 for the MNI coordinates and anatomical labels). The beta ERD was also highly significant at the medial aspect of temporal lobe and at the posterior cingulate/retrosplenial cortex that are closely interconnected and are thought to support memory retrieval (Vann, Aggleton, & Maguire, 2009). No spatial clusters on the lateral surface of the right hemisphere survived the correction for multiple comparisons.

**Table 2.** Brain regions showing the robust suppression of beta oscillations under verb generation task compared to the baseline (p < 0.05, Bonferoni-corr.). MNI coordinates and Brodmann areas (BA) are given for the most significant vertexes within the standard anatomical labels*.

| Brain region | BA | MNI |  |  |
| --- | --- | --- | --- | --- |
|  |  | x | y | z |
| *On the lateral surface* |  |  |  |  |
| L central sulcus | 4 | -45 | -14 | 30 |
| L precentral sulcus | 6 | -20 | -13 | 53 |
| L inferior frontal sulcus | 44 | -36 | 8 | 24 |
| L postcentral sulcus | 40 | -31 | -39 | 41 |
| L intraparietal sulcus | 7 | -27 | -51 | 36 |
| L superior temporal sulcus | 21 | -52 | -38 | -5 |
| L inferior temporal sulcus | 37 | -46 | -59 | -7 |
| L insula | 13 | -35 | 6 | 8 |
| *On the medial surface* |  |  |  |  |
| L superior frontal gyrus (supplementary motor area) | 6 | -7 | -16 | 53 |
| L paracentral lobule | 5 | -6 | -30 | 50 |
| L middle cingulate gyrus | 24 | -12 | -18 | 45 |
| L posterior cingulate gyrus dorsal | 23 | -4 | -34 | 28 |
| L posterior cingulate gyrus ventral (retrosplenial cortex) | 30 | -17 | -43 | 1 |
| L precuneus | 31 | -9 | -56 | 44 |
| L fusiform gyrus | 37 | -36 | -45 | -20 |
| L parahippocampal gyrus | 36 | -23 | -35 | -10 |
| R margin of the cingulate sulcus | 31 | 7 | -28 | 49 |
| R middle/posterior cingulate gyrus | 23 | 12 | -25 | 36 |
| R posterior cingulate gyrus dorsal | 23 | 6 | -34 | 32 |
| R precuneus | 31 | 7 | -45 | 42 |

\*based on the FreeSurfer aparc parcellation

### Strong Association vs Weak Association

#### Sensor-level analysis

To analyze the effect of verb retrieval demands on beta-band oscillations, we contrasted the beta ERD in SA and WA conditions using space-time SPM analysis. As expected, more difficult WA condition was accompanied by significantly greater beta suppression at approximately 700-550 ms before the response onset (p < 0.05, FWE-corr.). SA-WA difference in the beta ERD clustered around frontal midline sensors (Figure 2A)

To track the temporal dynamics of these difference, we reconstructed time courses of the beta ERD relatively to both stimulus and response onsets at the sensors representing the local maxima of the significant clusters. As shown in Figure 2B, the between-condition differences built up with time: while noun cue produced the comparable beta ERD under both conditions up to 500 ms post-stimulus, later on the beta suppression in WA trials reached greater values and lasted longer than in SA ones. The observed difference in beta-ERD strength between WA and SA conditions could be a mere artifact of the shift in response time that was an inevitable consequence of uneven SA-WA task difficulty. In order to refute this explanation, we performed an additional between-condition analysis on the peak values of the beta ERD at the maximal sensors (Figure 2C). The results were significant (Wilcoxon signed-rank test; p < 0.05; FDR-corr. for the number of the sensors), indicating that the greater beta ERD values in WA than in SA conditions could not be fully explained by the response time shift.

#### Source-level analysis

Based on the sensor-level results, we restricted MNE source reconstruction to the period −700 - −550 ms prior to the response onset. Figure 2D shows the cortical regions demonstrating statistically significant differences in beta suppression between SA and WA conditions (p < 0.05, uncorr.). The differential neural response involved cortical regions lying at the mesial aspects of frontal lobe bilaterally: anterior cingulate cortex, middle cingulate gyrus/sulcus with motor cingulate area, adjacent region of superior frontal gyrus comprising pre-SMA and SMA, as well as the entire precentral gyrus/sulcus on the lateral surface of the left hemisphere (Table 3).

**Table 3.** Brain regions showing significant differences in beta suppression between SA and WA conditions. MNI coordinates and Brodmann areas (BA) are given for the most significant vertexes within the standard anatomical labels*.

| Brain Region | BA | MNI |  |  |
| --- | --- | --- | --- | --- |
|  |  | x | y | z |
| *On the medial surface* |  |  |  |  |
| L anterior cingulate gyrus | 32 | -8 | 25 | 26 |
| L middle cingulate gyrus | 24 | -8 | -11 | 38 |
| L superior frontal gyrus (supplementary motor area) | 6 | -7 | 18 | 51 |
| R anterior cingulate sulcus | 32 | 7 | 44 | 25 |
| R middle cingulate gyrus | 24 | 9 | -15 | 38 |
| R superior frontal gyrus (supplementary motor area) | 6 | 7 | 18 | 50 |
| *On the lateral surface* |  |  |  |  |
| L precentral gyrus | 6 | -35 | 3 | 29 |
| L superior frontal gyrus | 8 | -17 | 40 | 38 |
| R middle frontal gyrus | 9 | 41 | 21 | 33 |

\*based on the FreeSurfer aparc parcellation

## Discussion

The aim of the present study was to test the hypothesis that verb retrieval from semantic memory engages the motor circuitry related to initiation and programming of motor acts and that this engagement plays a functional role in access to verb semantics. Therefore we expected that the more demanding memory search would recruit the motor system more heavily. We used suppression of MEG beta oscillations (15-30 Hz) to evaluate a degree of cortical activation related to motor system involvement in VG task while manipulating the difficulty of verb retrieval. Our participants were presented with the noun cues either strongly associated with a single verb (SA) that prompted the fast and effortless verb generation (mean RT = 1.22 sec), or weakly associated with multiple verbs (WA) that were more difficult to respond to (mean RT = 1.89 sec). We did not restricted the search for the differential activation to the specific motor ROI and performed a whole-brain analysis with rigorous statistics in anticipation that if our hypothesis was correct we would find the effect in the higher order prefrontal motor regions implicated in action initiation and planning.

Our finding shows that beta suppression starts 200-300 milliseconds after the noun cue presentation and then sustains until the overt generation of the appropriate verb regardless the response latency. Thus, the beta ERD spans the entire period dedicated to the search through semantic memory to obtain a target verb in VG task. In spatial terms the beta ERD during the search period is localized to a widely distributed left-hemispheric cortical network comprising the higher-order motor areas of the frontal lobes as well as classical auditory speech areas of temporal lobes and memory-related structures at the mesial temporal surface (Figure 1C, Table 2). Considering the involvement of the extensive language-related system, the beta ERD could reflect a co-activation of structures involved in recollection of the target verb. This fits well with the view that suppression of beta oscillations is linked to heightened sensorimotor transmission within any sensory domain not exclusively somatosensation (Engel & Fries, 2010; Kilavik, Zaepffel, Brovelli, MacKay, & Riehle, 2013). Within the context of verb generation task beta suppression might be related to both basic and higher-order processes operating across speech and motor domains.

Crucially, in accordance with our hypothesis, despite the spread of beta suppression over the whole left hemisphere, the differential effect of verb retrieval demands is confined to a set of cortical areas tightly related to initiating and planning of motor acts with a noticeable exception of primary motor cortex implicated in action execution (Figure 2D). A significant effect of task difficulty on motor system activation was observed in the −700 - −550 ms pre-response time window, which substantially precedes the preparation of vocal response and, most probably, overlaps with the stage of semantic search for the appropriate verb in the both conditions.

Since the greater beta ERD characterizes the more difficult task solved with the greater number of errors and the prolonged response latency, we cannot rule out the confounding domain-general factors related to difficulty/effort. For instance, greater attentional and/or memory resources allocated to the more difficult task could promote stronger brain activation (Dockstader, Cheyne, & Tannock, 2010; Pesonen, Hämäläinen, & Krause, 2007). However, had it been the case, the pattern of cortical activation indexing by beta suppression would have lacked motor specificity and would have comprised the entire large-scale network of the left hemisphere.

Another putative general factor accounting for the greater beta ERD in the pre-frontal areas during more difficult WA trials is the functional interaction between associative retrieval and executive control in VG task (Missier & Crescentini, 2011). As these authors proposed, in case of weakly associated noun-verb pairs the interference from the task-irrelevant responses, e.g. from strongly semantically associated nouns instead of the verbs, may require inhibitory processes, which are needed to suppress the erroneous word after retrieval. Such inhibition of the task-irrelevant word representations clearly relies upon frontal cortex capacities, as evidenced by the impaired interference control during the VG task in patients with focal frontal lesions (Thompson-Schill et al., 1998). However, findings from both frontal patients and healthy volunteers (Barch, Braver, Sabb, & Noll, 2000; Persson et al., 2004; Snyder, Banich, & Munakata, 2011; Thompson-Schill et al., 1997) indicate that the selection of the task-relevant verbs in VG task is mainly linked to the left inferior frontal gyrus (mid/posterior ventrolateral prefrontal cortex, VLPFC) as compared to other frontal regions. Thus, the executive control account is incompatible with the observed localization of differential activation in the mesial part of superior frontal gyrus comprising SMA and pre-SMA as well as in the middle cingulate cortex including cingulate motor area (Figure 2D).

Based on these considerations, the more plausible explanation of our findings is the one specifically implicating motor system into verb semantic retrieval. Effortful memory search for the appropriate verb requires greater allocation of processing resources in the higher-order motor areas than the easy task, in which the suitable verb is deftly retrieved due to the automatic nature of the underlying associative memory processes. Thus, our data provided rather strong evidence that the motor system contributes to the verb retrieval from semantic memory. However, it could still be debated what is the exact role of beta suppression in the mesial frontal motor areas and lateral premotor cortex in the access to verb meaning.

One potential account is the sensorimotor transformation that links speech processing with speech production during retaining a word in short-term memory in an active, readily available state (Cogan et al., 2014). In the process of laborious decision making occurring in difficult VG task, the search for the semantically and grammatically correct verb might be accompanied by erroneous retrieval of later rejected words (Missier & Crescentini, 2011), whose sub-vocal articulation activated the cortical premotor areas. Several convergent lines of evidence suggest that sensorimotor transformation cannot explain our data. First, brain activity related to sensorimotor transformation was shown to occupy both lateral areas of premotor cortex and the classical auditory language areas (Cogan et al., 2014; Piai, Roelofs, Rommers, & Maris, 2015), while SA-WA difference in beta suppression was restricted to higher-tier motor structures at the lateral and medial aspects of prefrontal cortex. Second, if caused by sensorimotor transformation, SA-WA differences would be solely related to duration of the beta ERD, whereas we observed both prolonged and greater beta ERD under WA condition (Figure 2B). Third, greater uncertainty in the anticipated vocal response was shown to attenuate the beta suppression in the core language-production areas of the left hemisphere (Piai et al., 2015), while in our experiment the difficulty of verb retrieval modulates the beta ERD in the opposite direction (Figure 2B).

We suggest that the prefrontal beta suppression in VG task could act to release task-relevant neural circuits to encode additional motor evidence needed for selecting the appropriate verb. Persistent activation of motor system during verb retrieval may be required for evidence accumulation as it was found for phoneme discrimination (D’Ausilio et al., 2009). Specifically, the beta ERD in higher-order motor areas of frontal cortex may represent a partial re-activation of motor circuitry involved in initiating and planning of motor sequence implying by verb meaning. Such motor re-activation may be redundant for verbs strongly associated with their nouns, since in this case fast activation spreading across the strong links of pre-formed purely linguistic associative network is sufficient for retrieving the intended verb synchronously with semantic processing of the presented noun (Butorina et al., 2017). After the failure of automatic verb retrieval, the following additional activation of shared motor representations may play a supportive role in the selection of the weakly associated verbs from semantic memory. On the other hand, a competition between multiple simultaneously re-activated motor plans may also elicit greater sensorimotor beta suppression, as it was previously shown for the sensory-motor task (Grent-’t-Jong, Oostenveld, Jensen, Medendorp, & Praamstra, 2013). Our finding agrees with both accounts that are not mutually exclusive as they both imply that action initiation and planning circuitry plays a role in verb retrieval from semantic memory.

In line with this, fMRI whole-brain analysis performed by Postle et al showed that pre-SMA and SMA were the only cortical areas demonstrating a selectivity for action meaning representations, as their heightened activation clearly dissociated both action words reading and action observations from all the other categories of linguistic stimuli and visual scenes (Postle et al., 2008). It may therefore be relevant that in our study the major verb-related differential activation in the beta band was coupled with mesial rather than lateral frontal cortical areas. It is noteworthy that mesial frontal motor cortical areas are also implicated in the initiation or changing a set of rules for selecting the motor programs (Nachev et al., 2008) that generally complies with their contribution in initiation and selection the motor associations for sequentially retrieved verbs in VG task. Further insight into the specificity of cortical architecture underlying verb semantics can be obtained in an MEG experiment, in which the participants are required to generated cue-related nouns or adjectives, conceivably revealing the unique contributions of the motor system into verb semantics.

## Conclusion

Our results show that the effortful extraction of the verbs from semantic memory accompanied by slower responses in verb generation task produces greater beta suppression confined to the higher-order motor areas of mesial and lateral pre-frontal cortex indicating contribution of motor system to an access to verbs’ representations. Since the pre-frontal motor areas are involved in maintaining abstract representations of actions during their initiating and planning, we argue that our finding provides a strong evidence for the proposed role of linguistic-motor association in retrieval of verb semantics.

## Acknowledgments

Work on this study was supported by the Russian Science Foundation (grant 14-28-00234) and Core Funding of MEG centre from Ministry of Science and Education of RF.

